# Integrated use of LC/MS/MS and LC/Q-TOF/MS targeted metabolomics with automated label-free microscopy for quantification of purine metabolites in cultured mammalian cells

**DOI:** 10.1101/490300

**Authors:** S. Eric Nybo, Jennifer T. Lamberts

**Author notes:** **Corresponding Author(*)** Jennifer T. Lamberts *Email:. **Funding information** This study was funded by a Ferris State University Exceptional Merit Grant to S.E.N. and an American Association of Colleges of Pharmacy New Investigator Award to J.T.L.

## Abstract

Purine metabolites have been implicated as clinically relevant biomarkers of worsening or improving Parkinson’s disease (PD) progression. However, the identification of purine molecules as biomarkers in PD has largely been determined using non-targeted metabolomics analysis. The primary goal of this study was to develop an economical targeted metabolomics approach for the routine detection of purine molecules in biological samples. Specifically, this project utilized LC/MS/MS and LC/QTOF/MS to accurately quantify levels of six purine molecules in samples from cultured N2a murine neuroblastoma cells. The targeted metabolomics workflow was integrated with automated label-free digital microscopy, which enabled normalization of purine concentration per unit cell in the absence of fluorescent dyes. The established method offered significantly enhanced selectivity compared to previously published procedures. In addition, this study demonstrates that a simple, quantitative targeted metabolomics approach can be developed to identify and quantify purine metabolites in biological samples. We envision that this method could be broadly applicable to quantification of purine metabolites from other complex biological samples, such as cerebrospinal fluid or blood.

## Introduction

Parkinson’s disease (PD) is characterized by gradual loss of dopaminergic neurons in the substantia nigra [1]. Presently, treatment options for PD are limited to replacement of dopamine with L-DOPA or treatments for delaying loss of motor function. Furthermore, the clinical assessment of PD progression is limited to the use of observational scales of motor dysfunction, such as the Unified Parkinson Disease Rating Scale (UPDRS), which often diagnoses the disease late in its progression [2]. Consequently, there is an urgent need for the identification of clinical biomarkers predictive of early-stage PD, including the use of nontargeted and targeted metabolomics analysis of biological specimens.

Traditionally, research into biomarkers for PD has focused on aggregation of alpha-synuclein protein or quantification of catecholamines in patient cerebrospinal fluid (CSF) [3, 4]. The detection of catecholamines in patient samples has limited utility for assessment of PD, in part because loss of dopamine is only reliably detected in patients who have not received drug treatment (i.e. drug-naïve patients) or who have discontinued L-DOPA therapy [5]. Recently, nontargeted metabolomics studies using ultra-high-performance liquid chromatography-quadrupole-time-of-flight high-resolution mass spectrometry (UPLC/QTOF/HRMS) analysis of patient samples has illuminated several classes of metabolites that are correlated with PD progression. Burte and coworkers screened serum samples of 41 early-stage PD patients for unique serological metabolites using nontargeted metabolomics screening [6]. From this study, Burte et al. identified 20 metabolites that were statistically-significant for prediction of PD progression and mild cognitive impairment. In another study, Lewitt and colleagues employed dual UPLC/MS/MS and GC-MS analysis of biofluids from PD patients to identify small-molecule biomarkers [7]. From the nontargeted metabolomics analysis, the authors identified 575 and 383 distinct biomarkers from plasma and CSF samples, respectively. Following multivariate analyses, the authors identified 15 unique biomarkers that were statistically correlated with PD progression, including three distinct purine molecules.

The purines are a family of ubiquitous biomolecules that play important roles in DNA synthesis and neurotransmission, among others. Purine molecules undergo a complex yet well-characterized metabolic degradation pathway (Figure 1). In the central nervous system (CNS), adenosine triphosphate (ATP, **1**) serves as an energy source and an extracellular signaling molecule that is released from secretory granules in response to an action potential [8]. **1** is metabolized by ectonucleoside triphosphate diphosphohydrolase (NTPDase) enzymes to form adenosine diphosphate (ADP, **2**) and adenosine monophosphate (AMP, **3**) [9]. **3** is metabolized to adenosine (**4**) by ecto-5’-nucleotidase [10]. Adenosine serves as a neuromodulator and has been shown to exhibit anti-inflammatory activity. **4** is taken up into the cell by a nucleoside transporter. Inside the cell, **4** is deaminated into inosine (**5**) by adenosine deaminase. In addition, **4** can be rapidly phosphorylated to **3** by adenosine kinase. Next, inosine is de-ribosylated by nucleoside phosphorylase to form hypoxanthine. Lastly, hypoxanthine is metabolized to uric acid (**6**) by xanthine oxidase [11].

**Fig 1.**
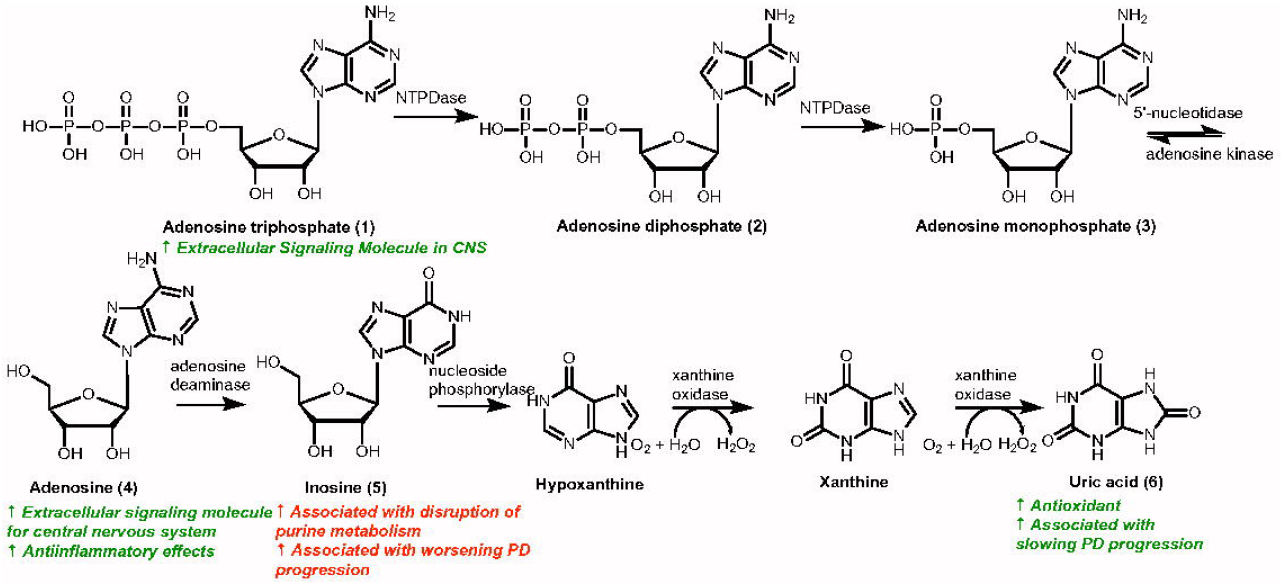
The purine nucleotide degradation pathway in the CNS. ATP (**1**) is an extracellular signaling molecule that is metabolized to ADP (**2**) and AMP (**3**) by various ectonucleosidases. **3** is metabolized by 5’-nucleotidase to form adenosine (**4**), which is taken up into the cell by nucleoside transporters. Adenosine (**4**) is a neuromodulator that exhibits anti-inflammatory effects. **4** is converted to inosine (**5**) via the action of adenosine deaminase. Accumulation of **5** is associated with worsening PD and disruption of the purine metabolic pathway. In addition, **4** can be rapidly phosphorylated to **3** by adenosine kinase. Nucleoside phosphorylase catalyzes de-ribosylation of **5** to hypoxanthine. Xanthine oxidase converts hypoxanthine to xanthine and uric acid (**6**), which is an antioxidant and is correlated with decreased PD progression

Purine metabolites have complex biological activities that have been shown to correlate with worsening or improving PD [5]. For instance, Lewitt et al. discovered that high concentrations of inosine were correlated with dysfunctional purine metabolism and PD progression [8]. Similarly, large quantities of ATP released from damaged cells can activate neuronal cell death pathways, causing neurodegeneration [12]. On the other hand, uric acid has been described as an important antioxidant and reactive oxygen species (ROS) scavenger that is correlated with slowed PD progression [13, 14]. Cortese et al. conducted a cohort study of Norwegian PD patients and discovered that high levels of urate were associated with lower overall risk of PD [15]. Furthermore, Miao and colleagues recently identified a genetic mutation in xanthine oxidase in PD patients that causes decreased uric acid synthesis [16]. Together, these findings suggest that some purine metabolites (i.e. ATP, inosine) correlate with neurodegeneration and worsening PD, while other metabolites (i.e. urate) are associated with decreased PD progression. However, the identification of purine molecules as biomarkers in PD was largely determined using non-targeted metabolomics analysis, and the relevance of these molecules to the pathophysiology of PD is still unknown. To answer this question, follow-on targeted metabolomics techniques must be developed for the routine measurement of purine biomarkers in various experimental settings. In this work, we developed an economical method for reliably measuring concentrations of purine levels using available instrumentation at the nearby Michigan State University Mass Spectrometry and Metabolomics Core.

From the outset, two questions were posed: first, could a quantitative, reproducible mass spectrometric method be developed to measure purine metabolites in cultured neuroblast cells, and secondly, which purines could be quantitated? To answer these questions, as a first step toward developing a targeted metabolomics approach for the routine detection of purine molecules in biological samples, this project utilized LC/MS/MS and LC/QTOF/MS to accurately quantify levels of six purine molecules in samples from cultured N2a murine neuroblastoma cells. Specifically, purine concentrations were measured in both conditioned media and whole cell lysates of undifferentiated (proliferating) and terminally-differentiated (post-mitotic) N2a cells. Differentiation of N2a cells by serum withdrawal produces a cellular phenotype that more closely resembles that of primary neurons [17]. This method provides significantly enhanced selectivity compared to previously published procedures [18, 19]. Moreover, mass spectrometry was integrated with automated label-free cellular microscopy to allow determination of purine content per cell without the use of fluorescent dyes or indirect biochemical measurements [20]. This study provides proof-of-concept for a procedure to identify and quantify purine metabolites in biological samples.

## Materials and methods

### Drugs and chemicals

Adenosine monophosphate (AMP, Catalogue #A1752-1G, Lot #SLBN6801V), adenosine diphosphate (ADP, Catalogue #A2754-1G, Lot #SLBL1267V), adenosine triphosphate (ATP, Catalogue #A6419-1G, Lot #SLBP3887V) and adenosine (Catalogue #A9251-5G, Lot #SLBL0630V) were all purchased from Sigma-Aldrich (St. Louis, MO, USA). Inosine (Catalogue #AC122250100, Lot #A0370995) and uric acid (Catalogue #AAA1334614, Lot #R19D020) were purchased from Thermo Fisher Scientific (Waltham, MA, USA). Methanol and water were of high-performance liquid chromatography grade while all purine reagents were of analytical grade.

Neuro-2a (N2a) murine neuroblastoma cells (Catalogue #CCL-131™) were purchased from ATCC^®^ (Manassas, VA, USA). Dulbecco’s Modified Eagle Medium (DMEM, Catalogue #10-569-010), Opti-MEM (Catalogue #22-600-050), 10X Trypsin-EDTA (Catalogue #15-400-050), Fetal Bovine Serum (FBS, Catalogue #SH3091003), and all other cell culture reagents were purchased from Thermo Fisher Scientific (Waltham, MA, USA).

### Cell culture conditions

N2a cells were maintained in DMEM supplemented with 10% FBS at 37°C under 5% CO_2_. For each experiment, cells were plated to 6-well plates in triplicate. Undifferentiated cells were plated at a ratio of 1:20 in DMEM + 10% FBS. Differentiated cells were plated at a ratio of 1:5 in OptiMEM without FBS. Plated cells were grown in a cell culture incubator at 37°C under 5% CO_2_ for 3, 4, or 6 days prior to label-free cell counting followed by sample extraction.

### Label-free cell counting

Total cell number per well was determined by performing label-free cell counting using a BioTek^®^ Cytation 1 Cell Imaging Multi-Mode Reader (BioTek Instruments, Inc., Winooski, VT, USA). Briefly, cells were removed from the cell culture incubator and transferred to the Cytation 1 instrument with the temperature control set to 37°C under room air. Cells were brought into focus with high-contrast brightfield imaging, and the focal height was then reduced by 250 μm until maximum contrast between cells and background was achieved. This “defocused” image was preprocessed using a rolling ball filter and analyzed for total cell count by thresholding, as previously described [19]. Defocused images were captured using high-contrast brightfield imaging with a 4x Olympus Plan Fluorite objective (N.A. 0.13) and the following exposure settings: LED, 5; Integration time, 12; Gain, 0. Montaging was used to generate a 19 x 16 image matrix that captured the majority (85%) of each well. Cell counts for each image were then summed to determine a total cell count for that well. The entire cell counting procedure took 30 min for one 6-well plate, following which samples were prepared for extraction.

### Sample extraction and preparation

The samples were separated into conditioned media and whole cell fractions for extraction and independent purine determination. For the media fraction, the residual cell culture media (2 mL) was pipetted from the plate and stored in a 15 mL conical tube. For the cell fraction, the attached cells were extracted with 1 mL of 80:20 (v/v) methanol:water and gently resuspended with a sterile spreader. The cells were transferred to a 4 mL glass extraction vial. The cells and media were stored at −80°C until time of analysis.

At the time of analysis, the media and cell fractions were thawed at room temperature. 1 mL of cell media was mixed with 1 mL of 80:20 (v/v) methanol:water in a 4 mL glass extraction vial and vortexed for 30 seconds. The cells were vortexed in the 80:20 (v/v) methanol:water for 30 seconds. The cells and media were then frozen in −80°C for 10 minutes and thawed at room temperature for 10 minutes before repeating the vortex procedure. The cells and media were subjected to 3 freeze and thaw cycles to ensure extraction of the purine analytes. The samples were then filtered using a syringe-driven nylon filter membrane (0.45 micron).

Three separate experiments were performed on different days with three biological replicates for each experimental condition (*n*=9). To maximize sample recovery, solid phase extraction was not performed, and matrix effect was determined to be negligible for the cultured mammalian cell samples (data not shown).

### Standard curve preparation

External standard curves for the purine metabolites analyzed in this study were prepared in a concentration range from 0.1 μg/mL to 10 μg/mL with four different concentrations of the working standards. The *r*^2^ values were ≥ 0.99 for each standard curve. Replicates were performed for each standard curve analysis, and the standard curves were analyzed alongside the experimental samples. The limits of detection (DLs) and quantification (QLs) were determined using the mathematical equations recommended by the International Conference on Harmonization (ICH) guidelines (ICH 2005).

### UPLC/MS/MS analysis of inosine, adenosine, and uric acid

UPLC/MS/MS tandem quadrupole analyses of adenosine (**4**), inosine (**5**), and uric acid (**6**) were conducted on a Waters Acquity TQD LC/MS/MS platform using 0.1% formic acid in H_2_O and 100% methanol as solvents and a reversed-phase column (Acquity^®^ UPLC HSS T3 column, 100 x 2.1 mm, 1.8 μm internal diameter). A linear gradient was used to separate the analytes: 1% to 25% methanol (0 to 4 minutes), 25% to 98% methanol (4 to 5 minutes), 98% methanol (5 to 7 minutes), 98% to 1% methanol (7 to 7.01 minutes), and re-equilibration of the column with 1% methanol (7.01 to 10 minutes) at a flow rate of 0.2 mL/min and a column temperature of 40.0°C. Experimental samples were analyzed alongside authenticated standards. The individual purine metabolites were detected in ESI+ and multiple reaction monitoring mode (MRM) using the following mass transitions: **4**: *m/z* 268 > 136; **5:** *m/z* 269 > 137; **6:** *m/z* 169 > 141.

### UPLC/QTOF/MS analysis of AMP, ADP, and ATP

UPLC/QTOF/MS analyses of ATP (**1**), ADP (**2**), and AMP (**3**) were carried out on a Waters Xevo G2-XS UPLC/MS/MS platform using 0.05% triethylamine (TEA) in H_2_O (pH = 7.0-8.0) (solvent A) and 30 mM ammonium acetate in 99:1 (v/v) methanol:H_2_O (solvent B) as solvents and a reversed-phase column (Acquity UPLC C_18_, 100 x 2.1 mm, 1.7 μm internal diameter). A linear gradient was used to separate the analytes: 0 to 20% solvent B (0 to 2 minutes), 20 to 50% solvent B (2 to 5 minutes), 50% solvent B (5 to 7 minutes), 50 to 0% solvent B (7 to 7.1 minutes), and re-equilibration of the column with 100% solvent A (7.1 to 10 minutes) at a flow rate of 0.3 mL/min and a column temperature of 40.0°C. Experimental samples were analyzed alongside authenticated standards. The individual purine nucleotide metabolites were detected in ESI-HRMS mode using enhanced target mass as follows: **1**: [M-H] = 505.9879±0.0005 *amu*; **2**: [M-H] = 426.0216±0.0005 *amu*; **3**: [M-H] = 346.0552±0.0005 *amu*.

### Statistical analysis

For conditioned media samples, analyte amounts were calculated in nanomolar. For whole cell lysates, total analyte mass per well was calculated in nanomoles and normalized to total cell count per well. Data were plotted using GraphPad Prism 6 data analysis software (GraphPad Software, La Jolla, CA, USA). Statistical analyses were conducted using ordinary one-factor ANOVA followed by Tukey’s post-hoc t-tests, where appropriate [20]. A *p* value less than 0.05 was considered significant, and a *p* value greater than or equal to 0.05 but less than 0.10 was considered a trend.

## Results

### Imaging of cultured N2a cells

High-contrast brightfield imaging of N2a mouse neuroblastoma cells in culture revealed qualitative differences in cell morphology following differentiation (Figure 2). In particular, cell bodies were smaller and more rounded after differentiating for three days, whereas undifferentiated cells adopted an elongated, fibroblast-like phenotype. In addition, differentiation resulted in an increase in the number of primary, axon-like processes and the quantity of branching points/arborization. Both morphologic changes are consistent with differentiated cells possessing a more neuron-like phenotype [17, 21]. Label-free cell counting using high-contrast brightfield imaging was effective in this setting given that cells were not exposed to molecular stains or dyes prior to sample extraction, as these compounds could affect cellular function and/or interfere with LC/MS analysis. Thus, the use of a multi-mode imaging reader allowed for rapid, automated cell counting in an integrated metabolomics workflow.

**Fig 2.**
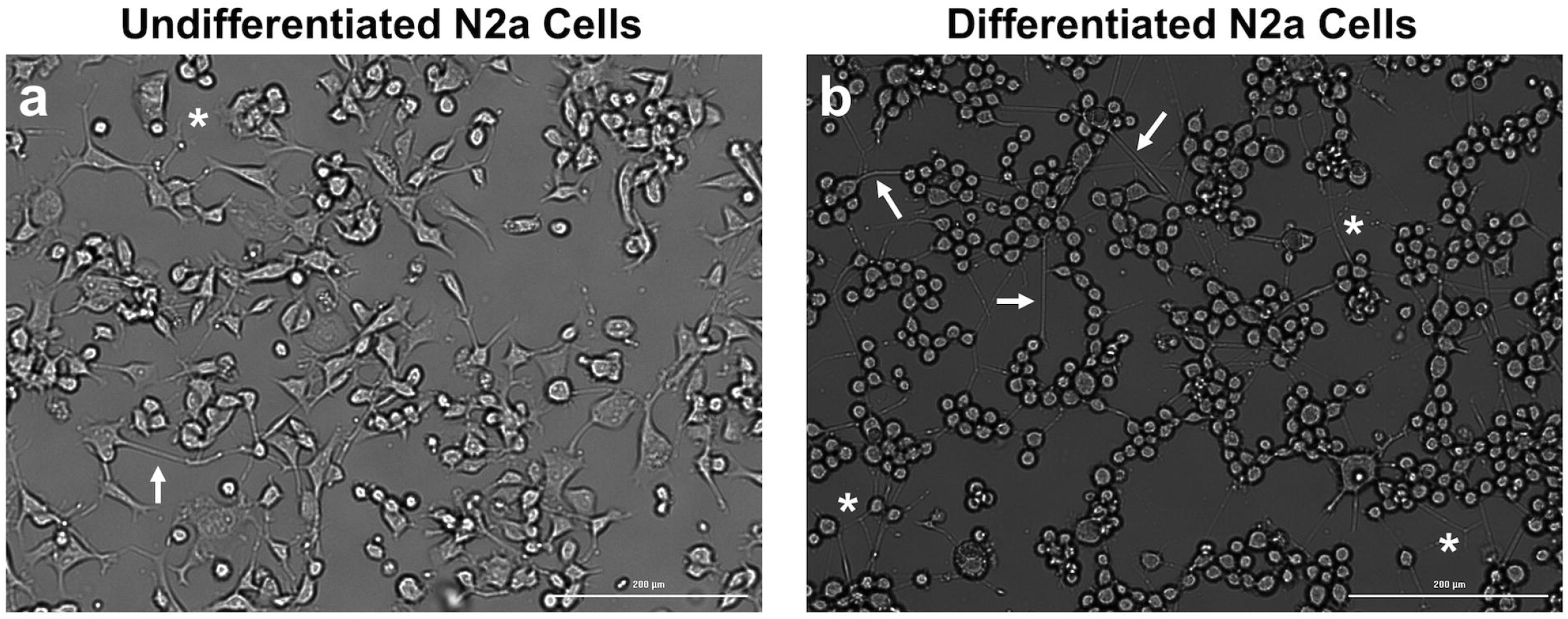
High-contrast brightfield imaging of N2a cells in culture. Defocused images of **a** undifferentiated and **b** differentiated N2a cells captured using a Cytation 1 Cell Imaging Multi-Mode Reader. Cells were plated to 6-well plates as described and grown in culture for 3 days prior to imaging. Images were captured using high-contrast brightfield imaging with a 10x Olympus Plan Fluorite objective (N.A. 0.3). Arrows denote primary, axon-like processes. Asterisks denote the presence of branching/arborization. Scale bar, 200 μm

### UPLC/QTOF/MS assay for quantification of 1, 2, and 3

To develop a mass spectrometric assay for quantification and unambiguous detection of ATP (**1**), ADP (**2**), and AMP (**3**), we prepared a standard curve of analytically pure standards (*r*^2^ ≥ 0.99) for development of a UPLC/QTOF/MS assay for separation and quantification of these metabolites. Initial attempts to separate ATP, ADP, and AMP via reversed-phase liquid chromatography were hindered by co-elution behavior of the purine nucleotides, which necessitated development of a method incorporating an ion pairing reagent. Incorporation of 30 mM ammonium acetate facilitated separation of **1-3** in a UPLC/QTOF/MS method that could detect the metabolites based on accurate mass (Supplementary Figures S1-S3). The lower limits of detection (LLOD) and quantification (LLOQ) for **1**, **2**, and **3** are reported in Table 1.

### UPLC/MS/MS assay for quantification of 4, 5, and 6

For the quantification of adenosine (**4**), inosine (**5**), and uric acid (**6**), a UPLC/MS/MS method was developed to unambiguously identify and quantify the metabolites. We prepared a standard curve of analytically pure **4**, **5**, and **6** for UPLC/MS/MS analysis (*r^2^* ≥ 0.99) and determined mass transitions to identify the metabolites in multiple reaction monitoring (MRM) mode. Based on this method, **4, 5,** and **6** were able to be identified and quantified (Figure 3). For **4** and **5**, the mass transitions were based on fragmentation of the ribose sugar and for **6** the mass transition was based on loss of H_2_O (Figure 4). The lower limits of detection (LLOD) and quantification (LLOQ) for **4**, **5**, and **6** are reported in Table 1.

**Fig 3.**
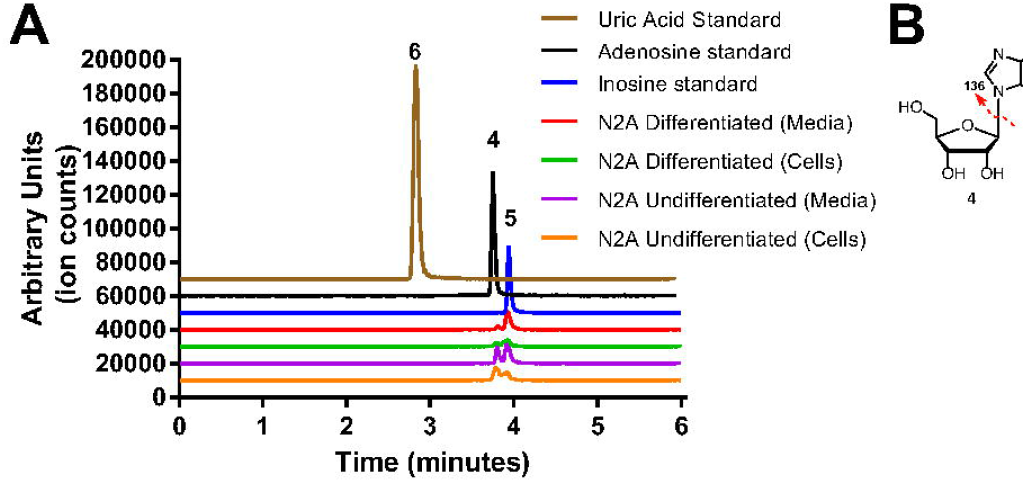
UPLC/ESI-/MS/MS chromatograms of 4, 5, and 6 and purines from N2a cells in MRM mode. **a** (From top to bottom) Chromatogram traces of (i) uric acid standard (**6,** *m/z* 169 > 141); (ii) adenosine standard (**4**, *m/z* 268 > 136); (iii) inosine standard (**5**, *m/z m/z* 269 > 137); representative chromatograms depicting purines from extracts of (iv) N2a differentiated cell media; (v) N2a differentiated cells; (vi) N2a undifferentiated cell media; and (vii) N2a undifferentiated cells. **b** Mass fragmentation patterns of transitions used to quantify **4, 5,** and **6** in MRM mode

**Fig 4.**
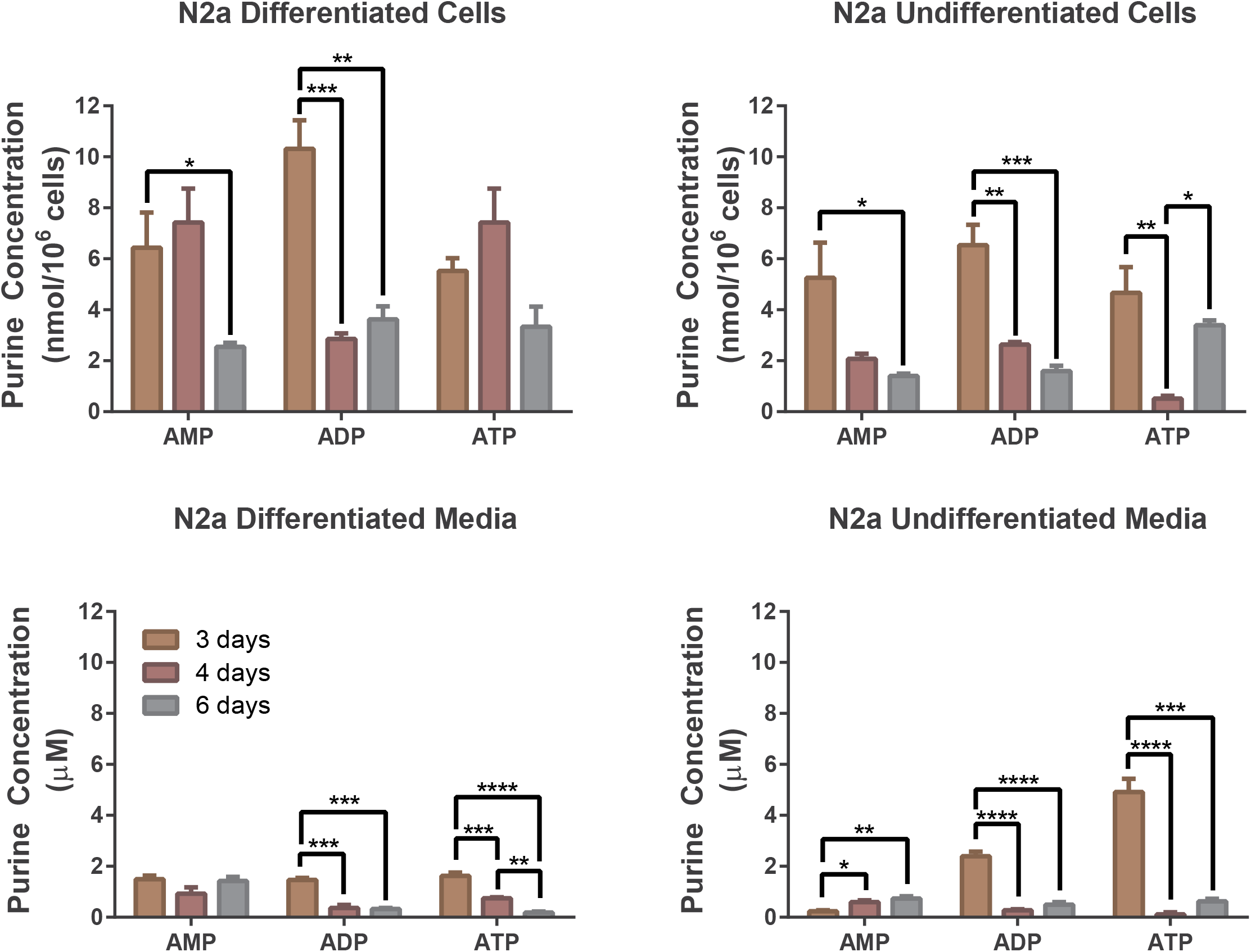
Concentrations of ATP, ADP, and AMP in cultured N2a cells on different days. Concentrations of **1**, **2**, and **3** were determined from N2a differentiated cells (upper left panel), N2a differentiated cell media (lower left panel), N2a undifferentiated cells (upper right panel), and N2a undifferentiated cell media (lower right panel). Nucleotide concentrations from cells were normalized to cell count (μmol/10^6^ cells). Extracts were generated on days 3, 4, and 6 (*n*=3). One-way ANOVA with Tukey’s post-hoc t-tests was conducted to determine statistical significance between experiments; \**p*<0.05, \*\**p*<0.01, \*\*\**p*<0.001, \*\*\*\**p*<0.0001

### Extraction and analysis of purines from cultured N2a cells

Detectable levels of ATP (**1**), ADP (**2**), AMP (**3**), adenosine (**4**), inosine (**5**), and uric acid (**6**) were observed in all biological samples. In general, the adenine nucleotides (**1-3**) were more abundant than the adenosine, inosine, or urate (**4-6**). Across all sample types, analyte levels were found to depend on the number of days in culture prior to sample extraction.

#### Adenine nucleotides

The adenine nucleotides ATP (**1**), ADP (**2**), and AMP (**3**) were detected in all samples analyzed (Figure 4). In all cases, the analytes in the samples were present in concentrations that far exceeded the LLOD established by the standard curves. In undifferentiated N2a cells, there was a significant decrease with time in the levels of **1** (F(2,6)=12.71, *p*=0.007), **2** (F(2,6)=29.21, *p*<0.001), and **3** (F(2,6)=6.56, *p*=0.031). In conditioned media from undifferentiated N2a cells, there was a significant reduction with time in the concentrations of **1** (F(2,6)=72.11, *p*<0.001) and **2** (F(2,6)=89.96, *p*<0.001). In contrast, the concentration of **3** (F(2,6)=13.31, *p*=0.006) increased with longer culture time in this sample.

A similar outcome was observed in samples from differentiated N2a cells. In lysates from differentiated N2a cells, **2** and **3** significantly decreased with time (**2**, F(2,6)=32.33, *p*<0.001; **3**, F(2,6)=5.40, *p*=0.046). Although there was not a significant change over time in the concentration of **1**, there was a trend toward decreasing amounts with increasing culture time (*p*=0.058). In media from differentiated N2a cells, there was a significant reduction in the concentrations of **1** (F(2,6)=70.85, *p*<0.001) and **2** (F(2,6)=53.3, *p*<0.001) over time, whereas the concentration of **3** did not change (*p*>0.10).

#### Adenosine, inosine, and urate

Adenosine (**4**), inosine (**5**), and urate (**6**) were detected in all samples tested in concentrations that were well above the LLOD established by the standard curves (Figure 5). There was a significant increase over time in the concentration of **5** detected in either undifferentiated N2a cells (F(2,6)=44.69, *p*<0.001) or differentiated N2a cells (F(2,6)=25.81, *p*=0.001). In contrast, concentrations of **4** and **6** did not change significantly with culture time in either sample type (*p*>0.10).

**Fig 5.**
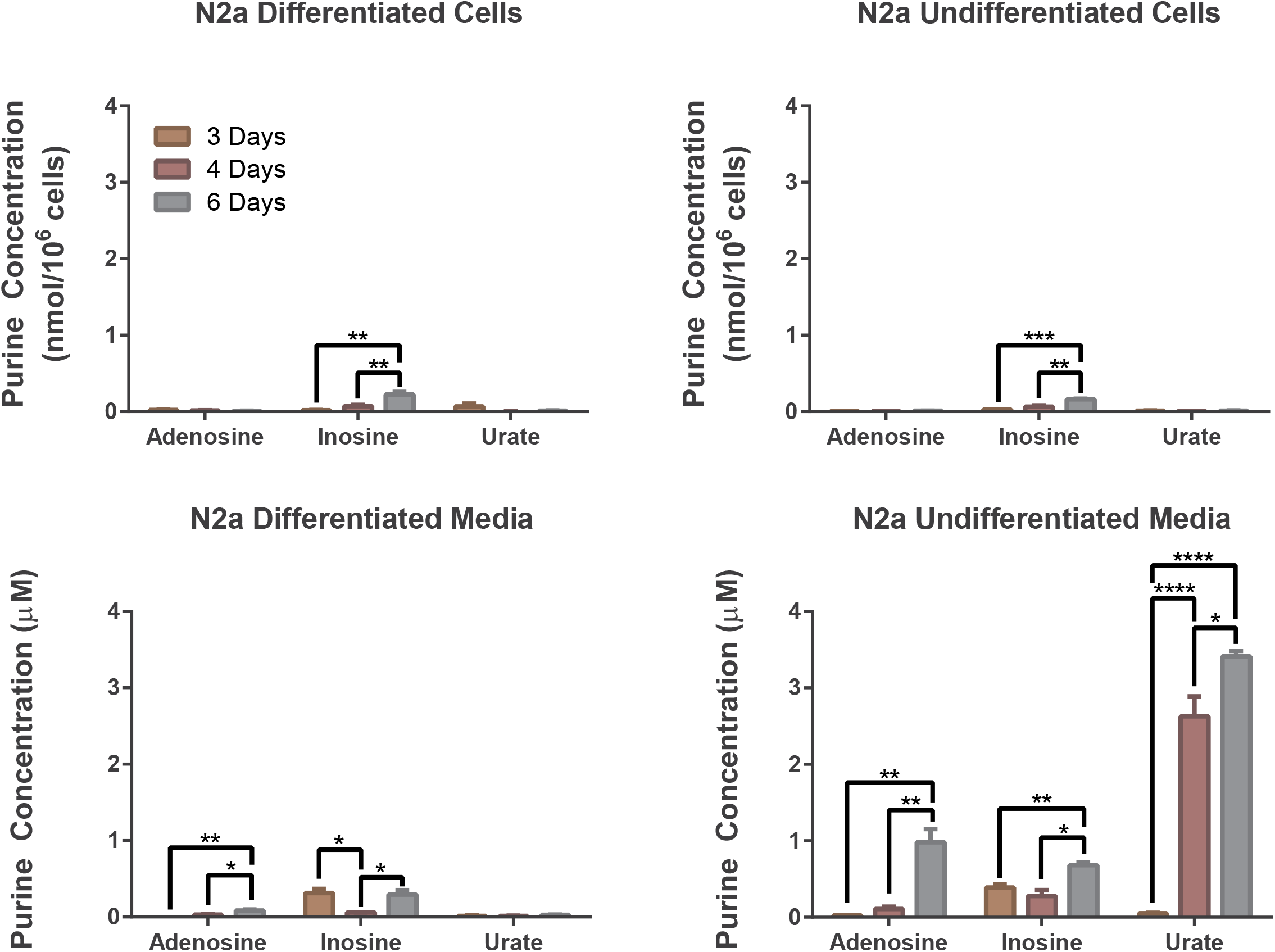
Concentrations of adenosine, inosine, and urate in cultured N2a cells on different days. Concentrations of **4**, **5**, and **6** were determined from N2a differentiated cells (upper left panel), N2a differentiated cell media (lower left panel), N2a undifferentiated cells (upper right panel), and N2a undifferentiated cell media (lower right panel). Nucleotide concentrations from cells were normalized to cell count (μmol/10^6^ cells). Extracts were generated on days 3, 4, and 6 (*n*=3). One-way ANOVA with Tukey’s post-hoc t-tests was conducted to determine statistical significance between experiments; \**p*<0.05, \*\**p*<0.01, \*\*\**p*<0.001, \*\*\*\**p*<0.0001

In media from undifferentiated N2a cells, the concentrations of **4** (F(2,6)=27.06, *p*=0.001), **5** (F(2,6)=14.61, *p*=0.005), and **6** (F(2,6)=125.1, *p*<0.001) significantly increased with time. In conditioned media from differentiated N2a cells, there was a significant increase with time in the levels of both **4** (F(2,6)=20.81, *p*=0.002) and **5** (F(2,6)=9.35, *p*=0.014). The concentration of **6** did not change significantly, although there was a trend toward increasing amounts with increasing culture time (*p*=0.056).

## Discussion

In this work, we developed a synergistic, reproducible targeted metabolomics method to detect and quantify purine metabolites in biologic samples. We developed a UPLC/QTOF/MS protocol for evaluation of nucleotide phosphates and a UPLC/MS/MS protocol for evaluation of purines. Specifically, we identified ATP (**1**), ADP (**2**), AMP (**3**), adenosine (**4**), inosine (**5**), and uric acid (**6**) in both cells and media from cultured N2a murine neuroblastoma cells. Furthermore, we evaluated the presence of these metabolites under two different experimental conditions over time: undifferentiated N2a cells and differentiated N2a cells. Targeted metabolomics was integrated with automated label-free cell counting to normalize purine concentration per number of cells, which enhanced comparison between different experimental groups. We envision that this method could be broadly applicable to quantification of all purine metabolites from other complex biological samples, such as cerebrospinal fluid or blood.

Both intracellular and extracellular adenine nucleotide concentrations decreased with increasing days in culture, with one exception: extracellular AMP either increased or stayed constant throughout the duration of the experiment. In general, this is consistent with decreasing cellular metabolic activity over time to meet decreasing nutrient availability, given that cell culture media was not replenished during the experiment. There are several possible explanations for the paradoxical increase in extracellular AMP. First, it is possible that AMP is being released from cells, which would reduce the intracellular pool while simultaneously increase the extracellular pool. Alternatively, these results could suggest that 5’-nucleotidase, the enzyme that metabolizes AMP, is either more abundant inside cells or possesses slow enzyme kinetics in this cell type.

The results of this study show that there exists relatively low concentrations of adenosine, inosine, and urate intracellularly, although inosine accumulated over time in both N2a cells and media. This result could suggest that nucleoside phosphorylase, the enzyme that metabolizes inosine, also possess slow enzyme kinetics in these cells. On the other hand, the concentrations of the adenosine, inosine, and urate were much greater in the extracellular environment. Specifically, extracellular adenosine increases over time. This could indicate either low activity of adenosine deaminase, or low cellular uptake of adenosine in this cell type. In addition, it was found that a large amount of urate builds up over time in media from undifferentiated but not differentiated N2a cells. In previous studies, it was shown that differentiation of N2a cells modifies purine metabolism [22]. The current study is in line with these findings and suggests that there are at least qualitative differences in purine metabolite concentrations after differentiating N2a cells. This difference between undifferentiated and differentiated cells underscores the importance of using clinically-relevant models for *in vitro* studies of purine metabolism, as differentiated N2a cells tend to more closely replicate the *in vivo* setting [17].

Previously, Aragon-Martinez et al. developed a liquid chromatographic method for quantification of ATP-related compounds in erythrocytes using a diode array detector [18]. In this method, the authors reported sensitivity for ATP-compounds with limits of detection from 0.001 to 0.097 *μ*mol/L and quantification limits from 0.004 to 0.294 *μ*mol/L [18]. Similarly, McFarland and coworkers developed a combined liquid chromatography UV-Vis assay for detection of adenosine, inosine, and hypoxanthine and detected urate and xanthine via coulometric detection at 150 and 450 mV [19]. Our method exhibited modest improvements in the lower limit of detection for ADP (6-fold improvement) and AMP (2-fold improvement) over the UV-Vis method developed by Aragon-Martinez et al. [18]. More importantly, our method features improved selectivity for ATP, ADP, and AMP by incorporating high resolution mass spectrometry to identify selected metabolites via enhanced target mass. In addition, our studies revealed that UPLC/MS/MS analysis in MRM mode enhances sensitivity for the detection of adenosine, inosine, and urate by 10-100-fold greater than other previously reported methods [18, 19].

The method developed in this study is significant for several reasons. First, the development of a UPLC/QTOF/MS analysis method for quantification of purine nucleotides provided a highly selective technique for identifying these molecules in a complex biological matrix. Second, the described techniques allow for assessment of a variety of structurally-diverse purine metabolites from the same experimental sample. Furthermore, this portends that the technique could be expanded to include the entire purine degradative pathway, including metabolites that would not be detected via UV-vis or enzyme assays (e.g. allantoin) [18]. Third, the incorporation of automated, label-free cellular microscopy added an additional quality inspection of mammalian cells to the workflow. High contrast brightfield imaging precludes the use of expensive and potentially bioactive reagents for measurement of cell numbers. For example, the use of cell stains could corrupt cellular metabolism and interfere with the downstream targeted metabolomics analysis.

The application of metabolomics techniques to experimental models of PD can afford key insights into the pathophysiology of neurodegenerative processes. For example, Lei and coworkers developed a neurotoxin model of PD to investigate changes in the energy/redox-metabolome of dopaminergic cells [23]. The authors used nuclear magnetic resonance (NMR) and direct-infusion electrospray ionization mass spectrometry (DI-ESI-MS) to identify changes in the metabolomic profile resulting from treatment with different environmental toxins. From these studies, the authors discovered that paraquat exposure resulted in increased levels of several metabolites from the pentose phosphate pathway, which was confirmed from subsequent ^13^C-glucose metabolic flux analysis. In addition, glucose-6-phosphate-dehydrogenase (G6PD) levels were increased as determined by proteomic analysis. Interestingly, the authors discovered that neurotoxin treatment decreased adenine nucleotide levels, possibly through depletion of pentose phosphate pathway metabolites. These studies provided a mechanism for paraquat-associated neurotoxicity, including disruption of NADPH metabolism, oxidative stress, and neuronal cell death. In a similar vein, assessment of the purine metabolites in the intracellular and extracellular environment conveys information concerning the energetics and redox state of the cells. This information can serve as a hallmark for aberrant cellular pathophysiology. Thus, the targeted metabolomics approach developed in this project serves as an important first step toward interrogating the role of the purine metabolome in PD pathophysiology. Given the central role of dopamine in PD pathogenesis, a logical next step for these studies is to quantify purine metabolites in N2a cells that have been differentiated into dopamine-producing neuronal-like cells [21]. The ultimate goal of this work is to quantify the purine metabolome in human samples as a means to assess improving or worsening PD progression.

## Supporting information

## Acknowledgements

LC/MS/MS targeted metabolomics experiments were performed at the Michigan State University Mass Spectrometry and Metabolomics Core Facility headed by Professor Dr. Dan Jones. The authors thank Dr. Anthony Schilmiller and Mrs. Lijun Chen for their help in conducting mass spectrometry analysis.

## Author contributions

All authors contributed to all aspects of experimentation and manuscript preparation.

## Compliance with ethical standards

### Ethical approval

This article does not contain any studies with human participants or animals performed by any of the authors.

## List of Abbreviations

ADP: Adenosine diphosphate
AMP: Adenosine monophosphate
ANOVA: Analysis of variance
ATP: Adenosine triphosphate
CNS: Central nervous system
CSF: Cerebrospinal fluid
DI-ESI-MS: Direct-infusion electrospray ionization mass spectrometry
DL: Detection limit
DMEM: Dulbecco’s Modified Eagle Medium
ESI: Electrospray ionization
FBS: Fetal bovine serum
G6PD: Glucose-6-phosphate-dehydrogenase
GC-MS: Gas chromatography-mass spectrometry
HRMS: High-resolution mass spectrometry
ICH: International Conference on Harmonisation
LC/MS/MS: Liquid chromatography-tandem mass spectrometry
LC/QTOF/MS: Liquid chromatography-quadrupole-time-of-flight-mass spectrometry
L-DOPA: L-3,4-dihydroxyphenylalanine
LLOD: Lower limit of detection
MRM: Multiple reaction monitoring
N2a: Neuro-2a
NADPH: Nicotinamide adenine dinucleotide phosphate
NMR: Nuclear magnetic resonance
NTPDase: Ectonucleoside triphosphate diphosphohydrolase
PD: Parkinson’s disease
QL: Quantification limit
QTOF/MS: Quadrupole-time-of-flight-mass spectrometry
UPDRS: Unified Parkinson’s Disease Rating Scale
UPLC: Ultra performance liquid chromatography
UV-vis: Ultraviolet-visible spectroscopy

## References

1. Poewe W, Seppi K, Tanner CM, et al (2017) Parkinson disease. Nat Rev Dis Prim 3:17013. https://doi.org/10.1038/nrdp.2017.13

2. Movement Disorder Society Task Force on Rating Scales for Parkinson’s Disease (2003) The Unified Parkinson’s Disease Rating Scale (UPDRS): status and recommendations. Mov Disord 18:738–50. https://doi.org/10.1002/mds.10473

3. Lamberts JT, Hildebrandt EN, Brundin P (2015) Spreading of α-synuclein in the face of axonal transport deficits in Parkinson’s disease: A speculative synthesis. Neurobiol. Dis. 77:276–283

4. Goldstein DS, Holmes C, Sharabi Y (2012) Cerebrospinal fluid biomarkers of central catecholamine deficiency in Parkinson’s disease and other synucleinopathies. Brain 135:1900–13. https://doi.org/10.1093/brain/aws055

5. Havelund JF, Heegaard NHHH, Færgeman NJKK, Gramsbergen JB (2017) Biomarker research in parkinson’s disease using metabolite profiling. Multidisciplinary Digital Publishing Institute

6. Burté F, Houghton D, Lowes H, et al (2017) metabolic profiling of Parkinson’s disease and mild cognitive impairment. Mov Disord 32:927–932. https://doi.org/10.1002/mds.26992

7. Lewitt PA, Li J, Lu M, et al (2017) Metabolomic biomarkers as strong correlates of Parkinson disease progression. Neurology 88:862–869. https://doi.org/10.1212/WNL.0000000000003663

8. Loeffler DA, LeWitt PA, Juneau PL, et al (1998) Altered guanosine and guanine concentrations in rabbit striatum following increased dopamine turnover. Brain Res Bull 45:297–9

9. Ipata PL, Balestri F, Camici M, Tozzi MG (2011) Molecular mechanisms of nucleoside recycling in the brain. Int J Biochem Cell Biol 43:140–5. https://doi.org/10.1016/j.biocel.2010.10.007

10. Zimmermann H, Zebisch M, Sträter N (2012) Cellular function and molecular structure of ecto-nucleotidases. Purinergic Signal 8:437–502. https://doi.org/10.1007/s11302-012-9309-4

11. Linden JM (1999) Purine Release and Metabolism. In: Siegel GJ, Agranoff BW, Albers RW (eds) Basic Neurochemistry: Molecular, Cellular and Medical Aspects. 6th edition. Lippincott-Raven, Philadelphia, pp 347–362

12. Burnstock G, Krügel U, Abbracchio MP, Illes P (2011) Purinergic signalling: From normal behaviour to pathological brain function. Prog Neurobiol 95:229–274. https://doi.org/10.1016/J.PNEUROBIO.2011.08.006

13. Ascherio A, LeWitt PA, Xu K, et al (2009) Urate as a predictor of the rate of clinical decline in Parkinson disease. Arch Neurol 66:1460–1468. https://doi.org/10.1001/archneurol.2009.247

14. Schwarzschild MA, Schwid SR, Marek K, et al (2008) Serum urate as a predictor of clinical and radiographic progression in Parkinson disease. Arch Neurol 65:716–23. https://doi.org/10.1001/archneur.2008.65.6.nct70003

15. Cortese M, Riise T, Engeland A, et al (2018) Urate and the risk of Parkinson’s disease in men and women. Park. Relat. Disord. 0

16. Miao V, Oise Coë Ffet-Legal M-F, Brian P, et al (2018) Daptomycin biosynthesis in Streptomyces roseosporus: cloning and analysis of the gene cluster and revision of peptide stereochemistry. Sat 27:57. https://doi.org/10.1099/mic.0.27757-0

17. Dickey CA, De Mesquita DD, Morgan D, Pennypacker KR (2004) Induction of memory-associated immediate early genes by nerve growth factor in rat primary cortical neurons and differentiated mouse Neuro2A cells. Neurosci Lett 366:10–4. https://doi.org/10.1016/j.neulet.2004.04.089

18. Aragon-Martinez OH, Galicia O, Isiordia-Espinoza MA, Martinez-Morales F (2014) A novel method for measuring the ATP-related compounds in human erythrocytes. Tohoku J Exp Med 233:205–14

19. McFarland NR, Burdett T, Desjardins CA, et al (2013) Postmortem brain levels of urate and precursors in Parkinson’s disease and related disorders. Neurodegener Dis 12:189–98. https://doi.org/10.1159/000346370

20. Drey LL, Graber MC, Bieschke J (2013) Counting unstained, confluent cells by modified bright-field microscopy. Biotechniques 55:28–33. https://doi.org/10.2144/000114056

21. Tremblay RG, Sikorska M, Sandhu JK, et al (2010) Differentiation of mouse Neuro 2A cells into dopamine neurons. J Neurosci Methods 186:60–7. https://doi.org/10.1016/j.jneumeth.2009.11.004

22. Gómez-Villafuertes R, Pintor J, Miras-Portugal MT, Gualix J (2014) Ectonucleotide pyrophosphatase/phosphodiesterase activity in Neuro-2a neuroblastoma cells: changes in expression associated with neuronal differentiation. J Neurochem 131:290–302. https://doi.org/10.1111/jnc.12794

23. Lei S, Zavala-Flores L, Garcia-Garcia A, et al (2014) Alterations in energy/redox metabolism induced by mitochondrial and environmental toxins: A specific role for glucose-6-phosphate-dehydrogenase and the pentose phosphate pathway in paraquat toxicity. ACS Chem Biol 9:2032–2048. https://doi.org/10.1021/cb400894a

