## Supplementary material for "Integrated use of LC/MS/MS and LC/Q-TOF/MS targeted metabolomics with automated label-free microscopy for quantification of purine metabolites in cultured mammalian cells"

**Electronic Supplementary Information**

*Purinergic Signaling*

**Supplementary Figure 1:** QTOF-ESI-MS chromatogram of ATP (1) standard in negative mode. ATP (1) was detected using enhanced target mass mode (505.9879 *amu*).

**Supplementary Figure 2:** QTOF-ESI-MS chromatogram of ADP (2) standard in negative mode. ADP (2) was detected using enhanced target mass mode (426.0216 *amu*).

**Supplementary Figure 3:** QTOF-ESI-MS chromatogram of AMP (3) standard in negative mode. AMP (3) was detected using enhanced target mass mode (346.0553 *amu*).

**Supplementary Figure 4:** QTOF-ESI-MS chromatogram of ADP (2) and ATP (3) in extracts of undifferentiated N2a cells in negative mode. ATP (1) and ADP (2) were detected using enhanced target mass mode (505.9879 *amu* and 426.0216 *amu* for **1** and **2**, respectively).

**Supplementary Figure 5:** QTOF-ESI-MS chromatogram of AMP (3) in extracts of undifferentiated N2a cells in negative mode. AMP (3) was detected using enhanced target mass mode (346.0553 *amu*).

**Supplementary Figure 6:** QTOF-ESI-MS chromatogram of ADP (2) and ATP (3) in extracts of differentiated N2a cells in negative mode. ATP (1) and ADP (2) were detected using enhanced target mass mode (505.9879 *amu* and 426.0216 *amu* for **1** and **2**, respectively).

**Supplementary Figure 7:** QTOF-ESI-MS chromatogram of AMP (3) in extracts of differentiated N2a cells in negative mode. AMP (3) was detected using enhanced target mass mode (346.0553 *amu*).

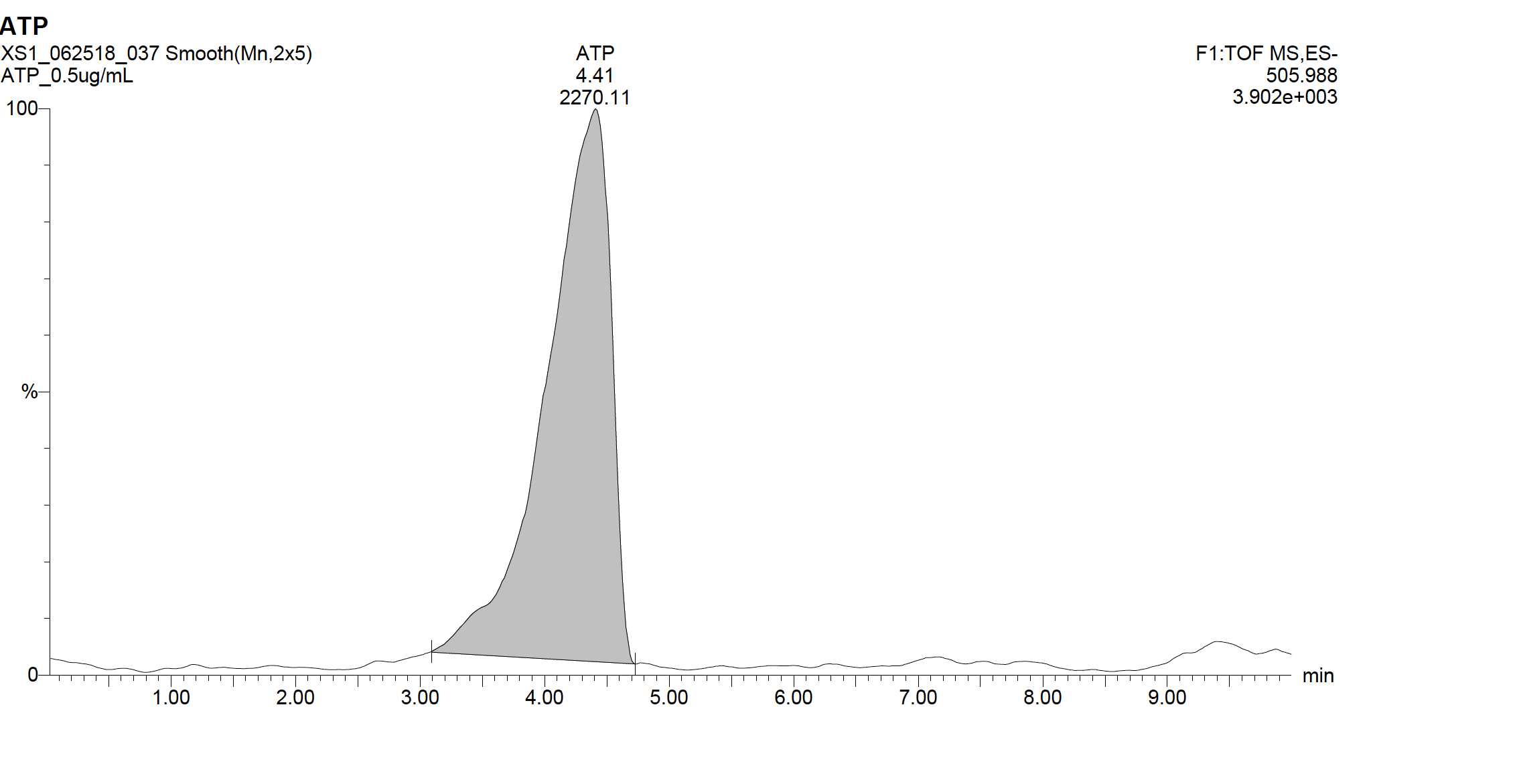

**Supplementary Figure 1:** QTOF-ESI-MS chromatogram of ATP (1) standard in negative mode. ATP (1) was detected using enhanced target mass mode (505.9879 *amu*).

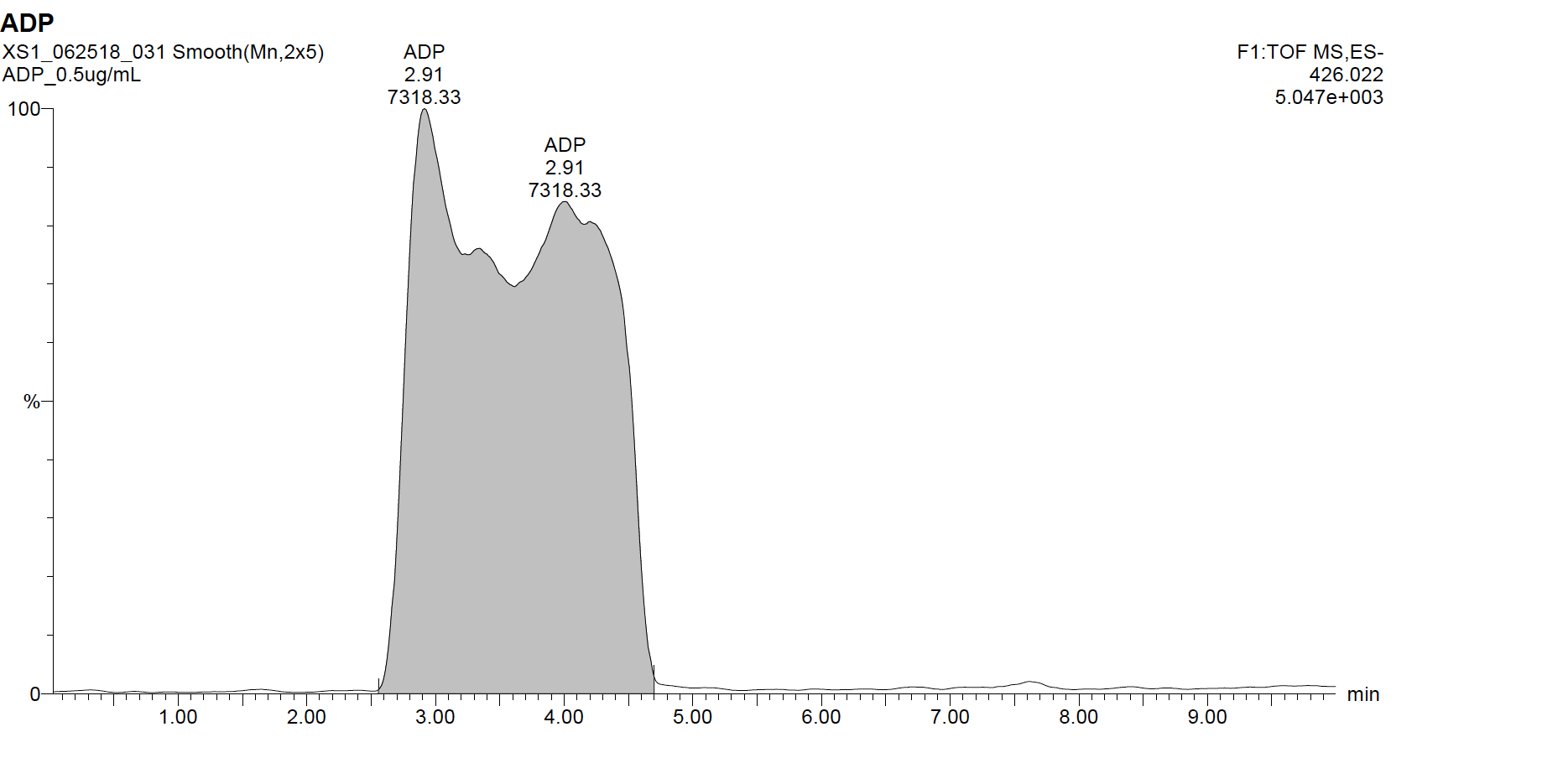

**Supplementary Figure 2:** QTOF-ESI-MS chromatogram of ADP (2) standard in negative mode. ADP (2) was detected using enhanced target mass mode (426.0216 *amu*).

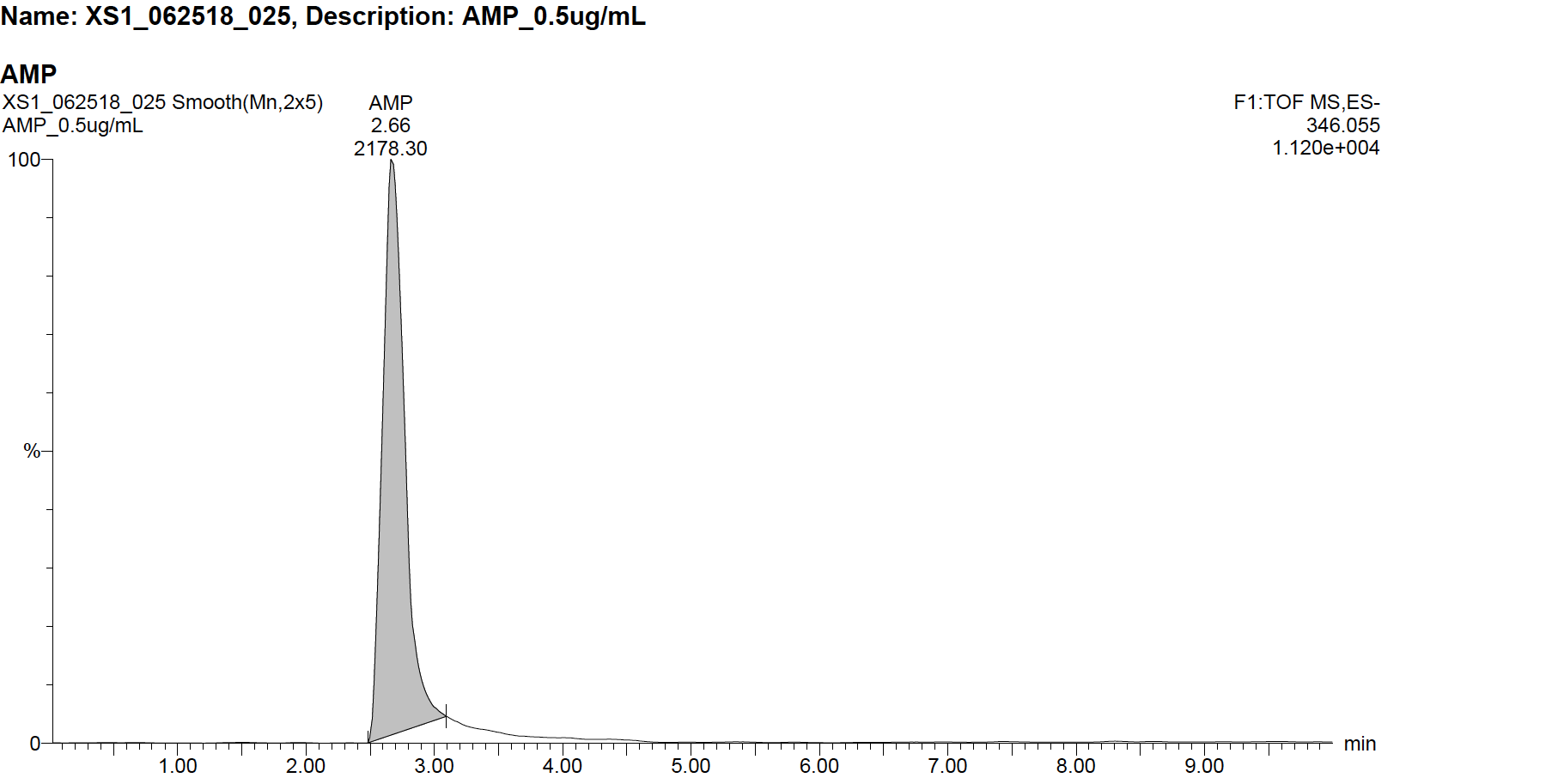
**Supplementary Figure 3:** QTOF-ESI-MS chromatogram of AMP (3) standard in negative mode. AMP (3) was detected using enhanced target mass mode (346.0553 *amu*).

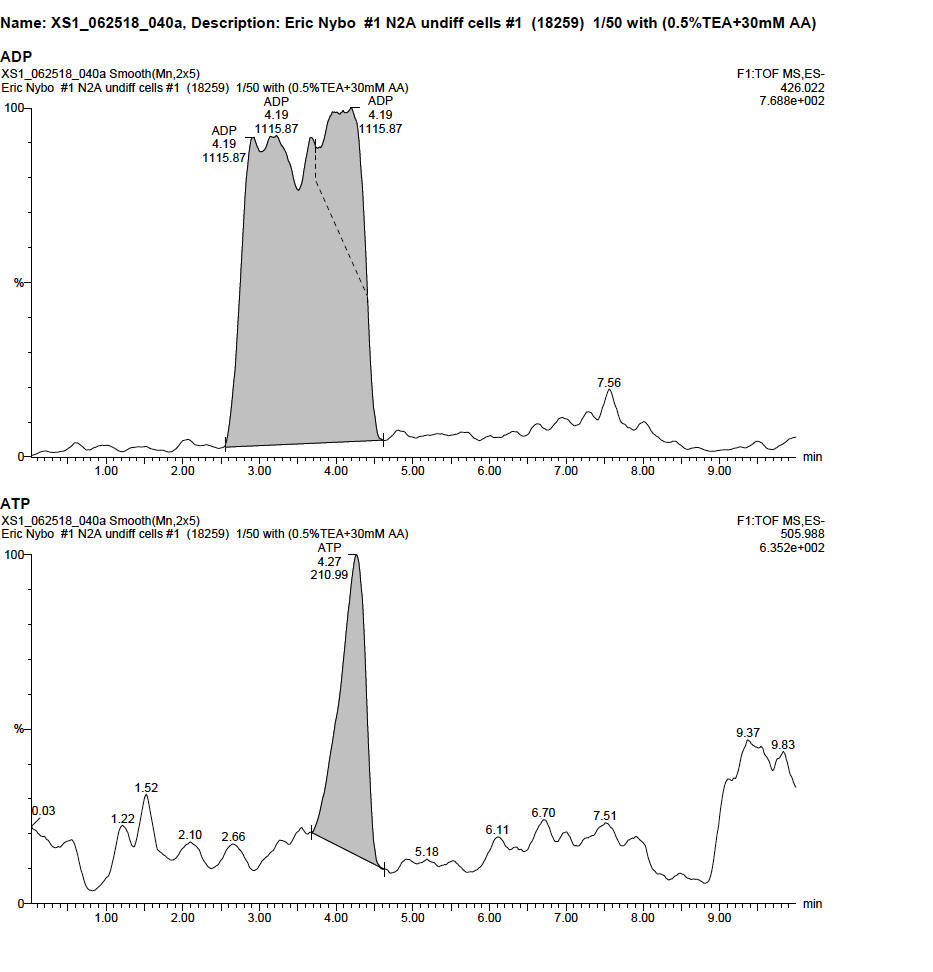

**Supplementary Figure 4** QTOF-ESI-MS chromatogram of ADP (2) and ATP (3) in extracts of undifferentiated N2a cells in negative mode. ATP (1) and ADP (2) were detected using enhanced target mass mode (505.9879 *amu* and 426.0216 *amu* for **1** and **2**, respectively).

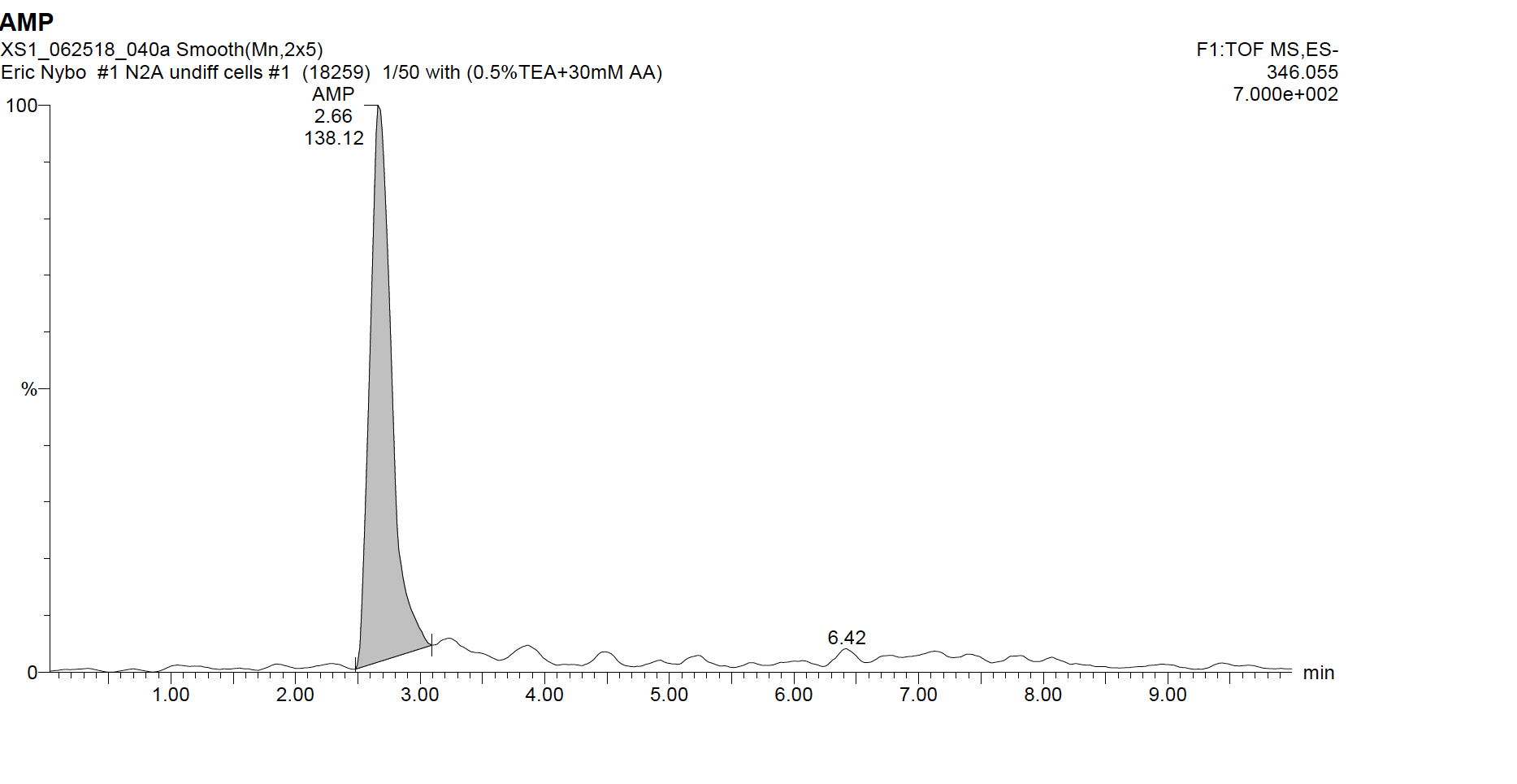

**Supplementary Figure 5** QTOF-ESI-MS chromatogram of AMP (3) in extracts of undifferentiated N2a cells in negative mode. AMP (3) was detected using enhanced target mass mode (346.0553 *amu*).

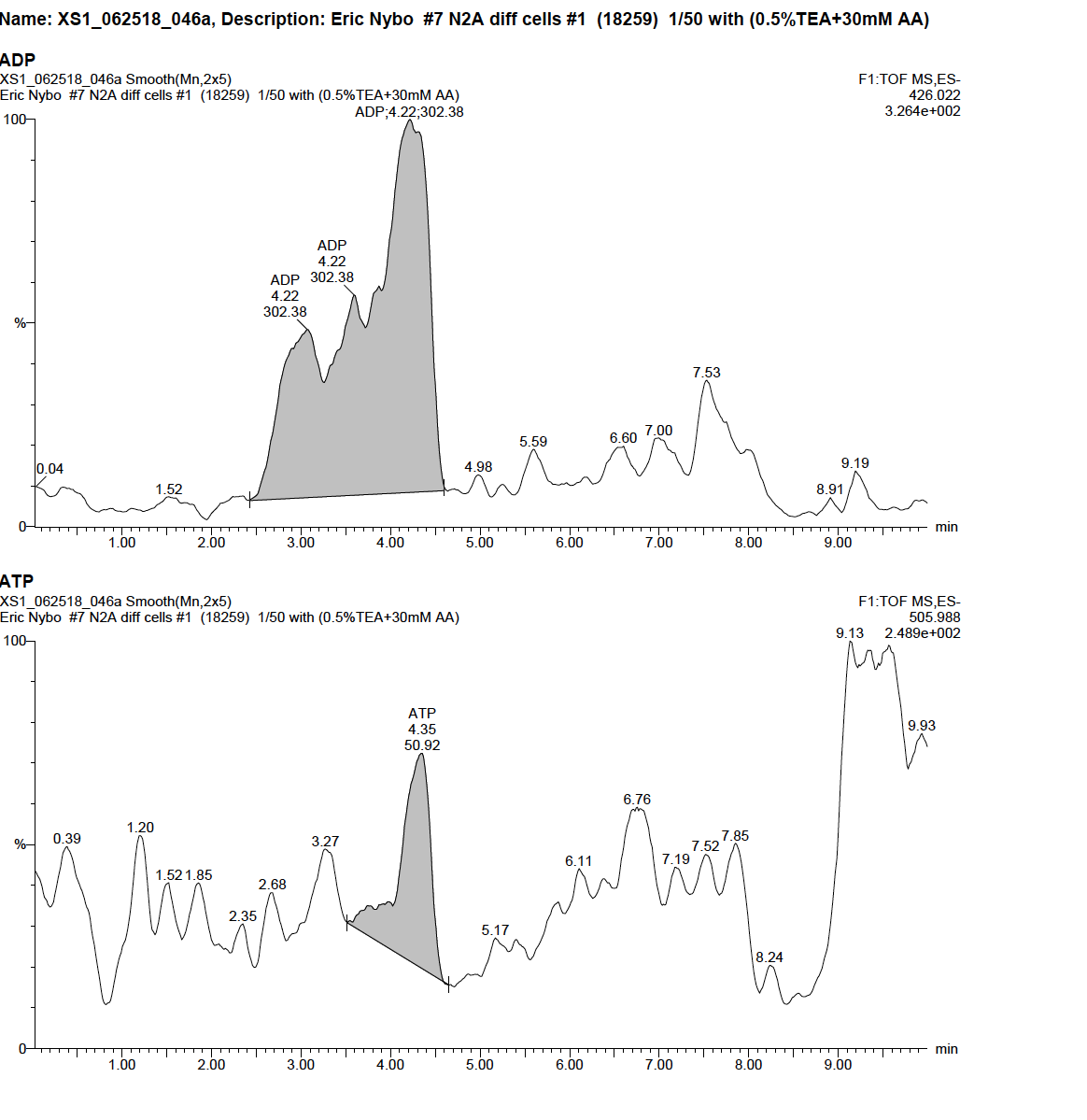

**Supplementary Figure 6** QTOF-ESI-MS chromatogram of ADP (2) and ATP (3) in extracts of differentiated N2a cells in negative mode. ATP (1) and ADP (2) were detected using enhanced target mass mode (505.9879 *amu* and 426.0216 *amu* for **1** and **2**, respectively).

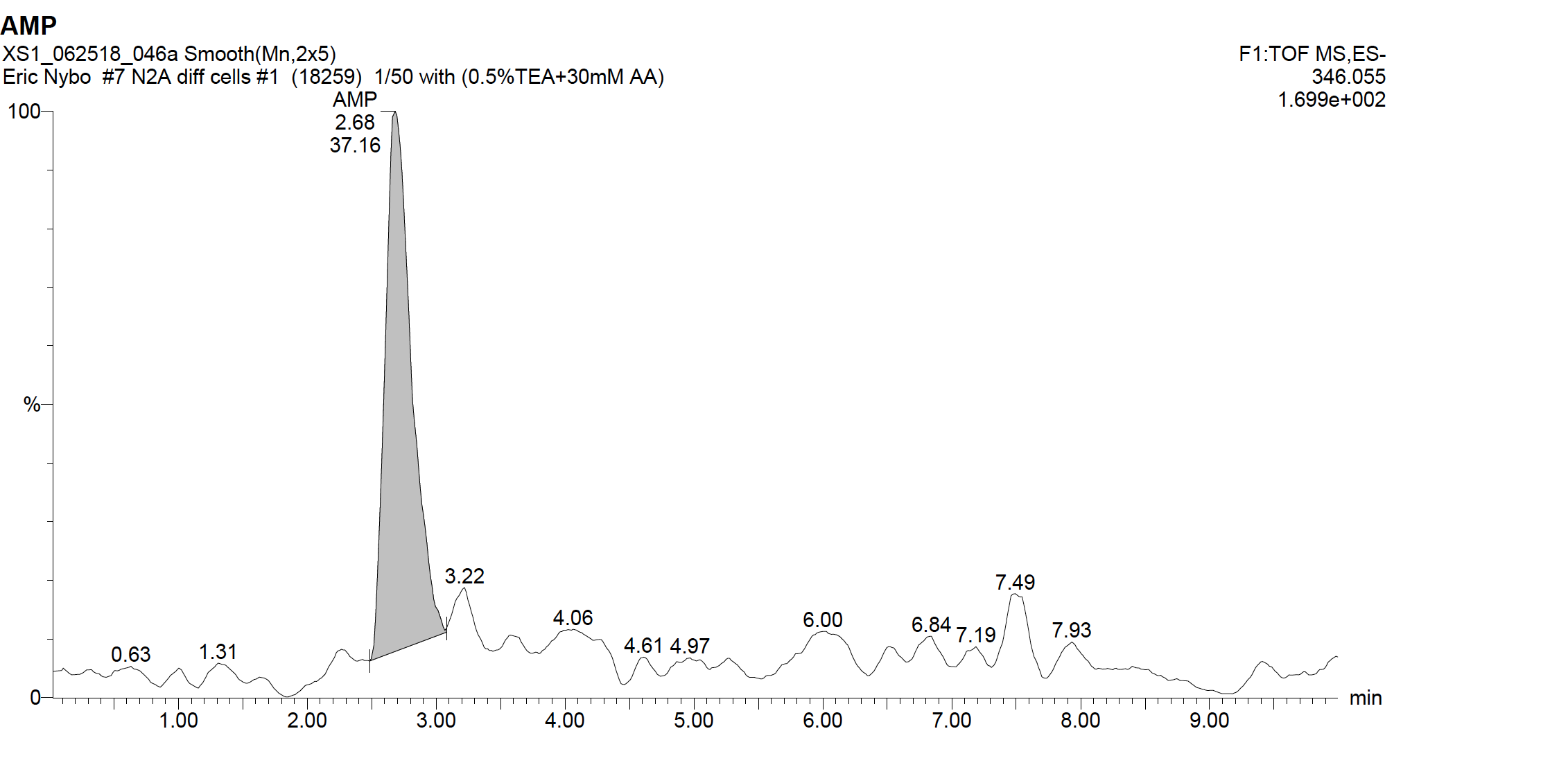

**Supplementary Figure 7** QTOF-ESI-MS chromatogram of AMP (3) in extracts of differentiated N2a cells in negative mode. AMP (3) was detected using enhanced target mass mode (346.055 *amu*).
